# Uncovering and mitigating bias in large, automated MRI analyses of brain development

**DOI:** 10.1101/2023.02.28.530498

**Authors:** Safia Elyounssi, Keiko Kunitoki, Jacqueline A. Clauss, Eline Laurent, Kristina Kane, Dylan E. Hughes, Casey E. Hopkinson, Oren Bazer, Rachel Freed Sussman, Alysa E. Doyle, Hang Lee, Brenden Tervo-Clemmens, Hamdi Eryilmaz, Randy L. Gollub, Deanna M. Barch, Theodore D. Satterthwaite, Kevin F. Dowling, Joshua L. Roffman

## Abstract

Large, population-based MRI studies of adolescents promise transformational insights into neurodevelopment and mental illness risk ^1, 2^. However, MRI studies of youth are especially susceptible to motion and other artifacts ^3, 4^. These artifacts may go undetected by automated quality control (QC) methods that are preferred in high-throughput imaging studies, ^5^ and can potentially introduce non-random noise into clinical association analyses. Here we demonstrate bias in structural MRI analyses of children due to inclusion of lower quality images, as identified through rigorous visual quality control of 11,263 T1 MRI scans obtained at age 9-10 through the Adolescent Brain Cognitive Development (ABCD) Study^6^. Compared to the best-rated images (44.9% of the sample), lower-quality images generally associated with decreased cortical thickness and increased cortical surface area measures (Cohen’s d 0.14-2.84). Variable image quality led to counterintuitive patterns in analyses that associated structural MRI and clinical measures, as inclusion of lower-quality scans altered apparent effect sizes in ways that increased risk for both false positives and negatives. Quality-related biases were partially mitigated by controlling for surface hole number, an automated index of topological complexity that differentiated lower-quality scans with good specificity at Baseline (0.81-0.93) and in 1,000 Year 2 scans (0.88-1.00). However, even among the highest-rated images, subtle topological errors occurred during image preprocessing, and their correction through manual edits significantly and reproducibly changed thickness measurements across much of the cortex (d 0.15-0.92). These findings demonstrate that inadequate QC of youth structural MRI scans can undermine advantages of large sample size to detect meaningful associations.

## Introduction

Magnetic resonance imaging (MRI) is widely used in clinical neuroscience research to study neuroanatomical variation in healthy individuals as well as those with neuropsychiatric disease ^7^. Structural (T1-weighted) MRI scans (sMRI) provide reliable, individual-level indices of cortical thickness, surface area, and volume, and enable registration of other brain imaging data (such as functional MRI and PET) to anatomical templates that facilitate group-level analyses ^8^. In accordance with neurodevelopmental models of mental illness, large-scale brain MRI studies of children and adolescents offer potential to elaborate neural signatures of emergent psychopathology ^1, 2^. Such insights could be harnessed in efforts to develop improved early recognition and treatment, outcomes that may help ameliorate the current youth mental health crisis ^9^. As such, the US National Institutes of Health and other funding agencies have invested heavily in longitudinal MRI studies of adolescent brain development, such as the ongoing ABCD Study ^6^.

Recent work has underscored the need for thousands of participants in such clinical MRI studies, as within-group variation is considerable and effect sizes for relationships between psychopathology and MRI indices tend to be small ^2^. Further, MRI scans of children and adolescents are particularly susceptible to artifact due to participant motion within the scanner ^3, 4^. An unanswered question concerns whether large sample size – e.g. in studies involving thousands of participants – sufficiently compensates for errant sMRI measurements arising from inclusion of poorer quality images. Alternatively, smaller studies have suggested the possibility that visible motion artifact results not only in random noise but in bias ^3, 4^, which (again) may or may not be sufficiently offset by inclusion of more participants.

A related question concerns the adequacy of automated quality control (QC) measures, applied during scan acquisition, processing, or analysis, to identify or adjust for poor quality images in large sMRI studies of children. Notably, unlike for functional MRI, head motion is less routinely quantified as part of sMRI analyses, and its effects on sMRI measurements have been less well studied – although some prior work has associated induced or measured motion with bias in sMRI estimates ^3, 10^. Newer sMRI sequences, including those deployed on Siemens and GE magnets in ABCD ^11, 12^, have incorporated real-time motion correction protocols that re-acquire data immediately after significant motion is detected. Whether this feature mitigates artifact sufficiently well to prevent bias in large-scale studies remains uncertain. Image preprocessing software can provide automated QC metrics, such as the overall “pass/fail” rating in the FreeSurfer processing stream. ^13^ This metric is used by ABCD in conjunction with raw data screens and clinical (radiology) evaluations to provide an overall recommendation on whether to include images in analyses. However, routine automated QC measures have shown inconsistent sensitivity to detect artifact identified by manual (visual) QC ratings of sMRI scans in youth^5, 14^.

As such, a final consideration – one especially pertinent to large-scale studies such as ABCD, which is collecting 6 sets of MRI scans over 10 years from >10,000 youth participants – is the added value of manual QC of postprocessed sMRI scans, and of the even more time- and resource-intensive process of manual cortical edits^15, 16^, to minimize artifact-related errors. Depending on image quality, manual edits of a single scan can take a skilled technician as few as 30 minutes to as long as several days to complete. While the utility of manual edits in identifying case-control differences in pediatric sMRI studies has been questioned ^17–19^, their importance to accurately detecting subtle neurodevelopmental differences among youth is evident in other studies^20, 21^.

Here we conducted in-depth, manual QC assessments of >12,000 sMRI images obtained at Baseline (age 9-10) and Year 2 follow-up (age 11-12) from ABCD study participants. We then characterized the impact of poorer-quality scans on the fidelity of sMRI measurements (cortical thickness, surface area, and volume) and the on reliability sMRI-clinical associations. Further, in light of efficiency considerations, we evaluated the sensitivity of automated QC to detect poorer-quality scans, and contrasted the effectiveness and reliability of several automated and manual error mitigation strategies that varied by labor intensity.

## Results

### Manual quality control (MQC) ratings in Baseline scans

The ABCD study enrolled 11,875 participants, age 9 or 10 at Baseline, across 22 U.S. sites. Participant race and ethnicity mirrored those of the U.S., and enrollment was enriched for multiple births and siblings from multiple pregnancies ^22^. Structural MRI (sMRI) scans were obtained from participants on 3T Siemens, Philips, or GE magnets as described in **Methods** and by Casey and colleagues.^23^ Minimally processed T1 volumes were available from the NIMH Data Archive (NDA) for all but 160 participants. After removing those marked as requiring clinical consultation, T1 volumes for the remaining scans were downloaded from the NDA and pre-processed in FreeSurfer version 7.1. While several processing streams are available to process and analyze sMRI data, the present analyses used FreeSurfer software for two reasons: first, existing, tabulated region-of-interest sMRI analyses available through the NIMH Data Archive and widely used in published ABCD analyses were conducted wither FreeSurfer; and second, FreeSurfer offers manual cortical edit capabilities. Following training and calibration (**Methods**), a single research coordinator (S.E.) who was blind to subject-level information then viewed each MRI volume individually. This approach was chosen because it eliminated concerns over inter-rater reliability, which has previously been shown to be modest (∼0.75) when including multiple tiers of sMRI QC,^5^ but could alternatively be assessed for intra-rater reliability (e.g., drift in ratings over time) and for triangulation with automated QC measures. During manual review an additional 740 scans were removed from further consideration due to the presence of cysts >1 cm^3^, and 228 were omitted from the main analyses due to segmentation errors and related signal dropout that persisted after a second round of preprocessing (**Figure 1a**).

**Figure 1.**
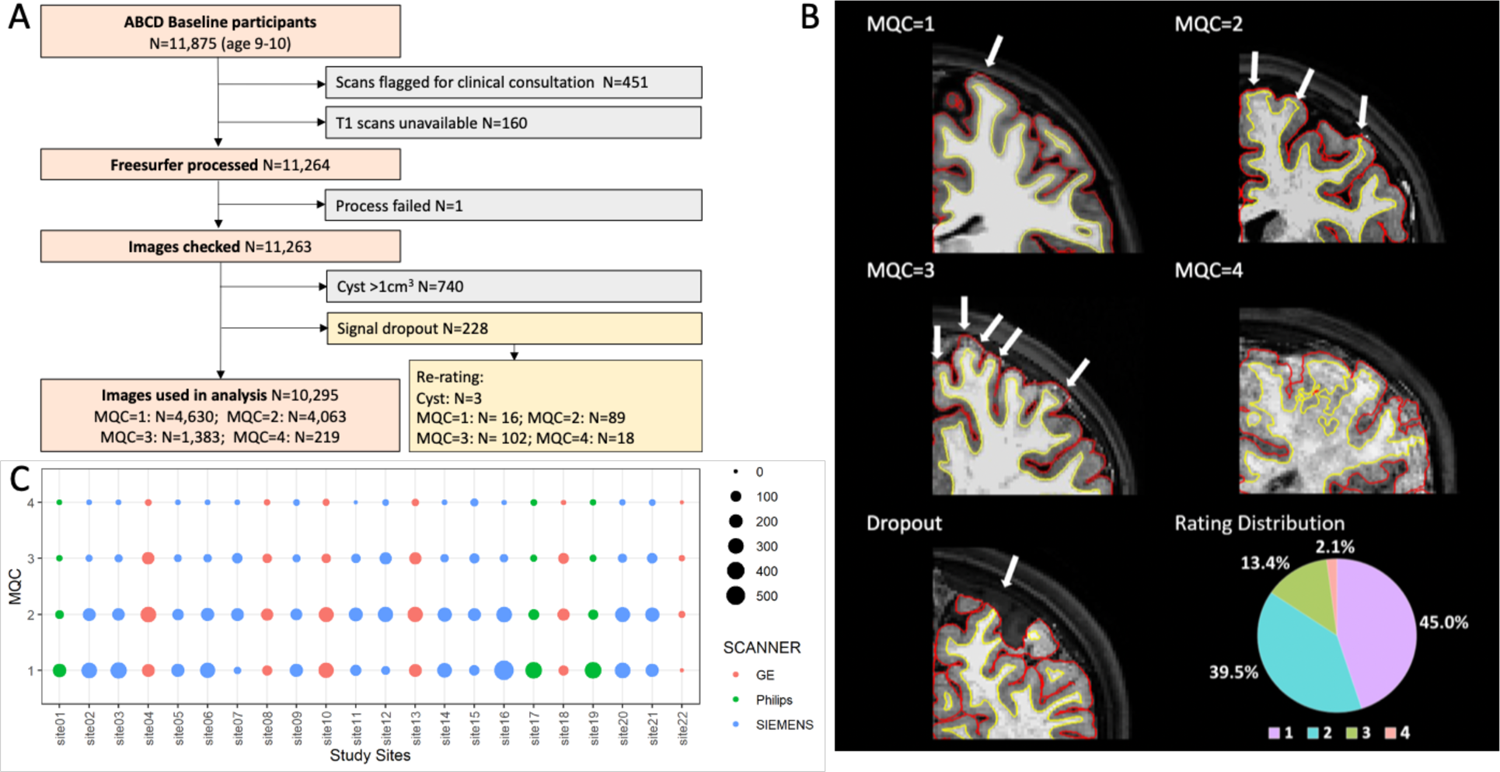
Manual quality control (MQC) protocol. (A) Among 11,875 total participants at Baseline, we excluded participants with clinical findings (see Methods), broken or blank T1 images, or repeated failed FreeSurfer preprocessing. After excluding additional images found to have cysts or signal dropout, we rated 10,295 images on MQC=1-4 scale (1: best, 4: worst). (B) Distribution of MQC ratings, stratified by site and scanner. (C) Representative examples of MQC=1-4 scans and a scan with apparent tissue loss due to segmentation error. Arrows indicate areas where manual edits are needed to correct for errant automated segmentation of the pial surface from the underlying cortex. Distribution of MQC ratings among all scans is displayed at lower right.

The remaining 10,295 T1 scans received manual quality control (MQC) ratings. Ratings were based on overall appearance of the entire T1 volumes, as follows: “1” (requiring minimal edits, n=4,630, 45.0%), “2” (requiring moderate edits, n=4,063, 39.5%), “3” (requiring substantial edits, n=1,383, 13.4%), or “4” (unusable, n=219, 2.1%) (**Figure 1b,c**). We have uploaded these MQC ratings to the NDA (see **Data Availability**). Demographic, clinical, and scanner characteristics of participants stratified by MQC group are described in **Table S1a**. Individuals with higher quality scans tended to be slightly older and female, also demonstrated less externalizing psychopathology and total symptoms on the Child Behavior Checklist (CBCL). Scan quality also differed by scanner manufacturer; notably, the mean MQC rating for images from Philips magnets (1.34, 95% CI 1.29-1.38), which were not subject to real-time motion correction, was more favorable than those for Siemens (1.71, 95% CI 1.69-1.73) and GE (1.96, 95% CI 1.93-1.99), which did include this feature (p’s<.0001, after controlling for age, gender, and psychopathology). MQC ratings were stable over the sequence of scan evaluations after controlling for each of these factors (see **Extended Data Figure 1a, Table S2**), and their distribution did not change in sensitivity analyses that included the 228 scans with segmentation errors (**Extended Data Figure 2, Table S1b**).

**Figure 2.**
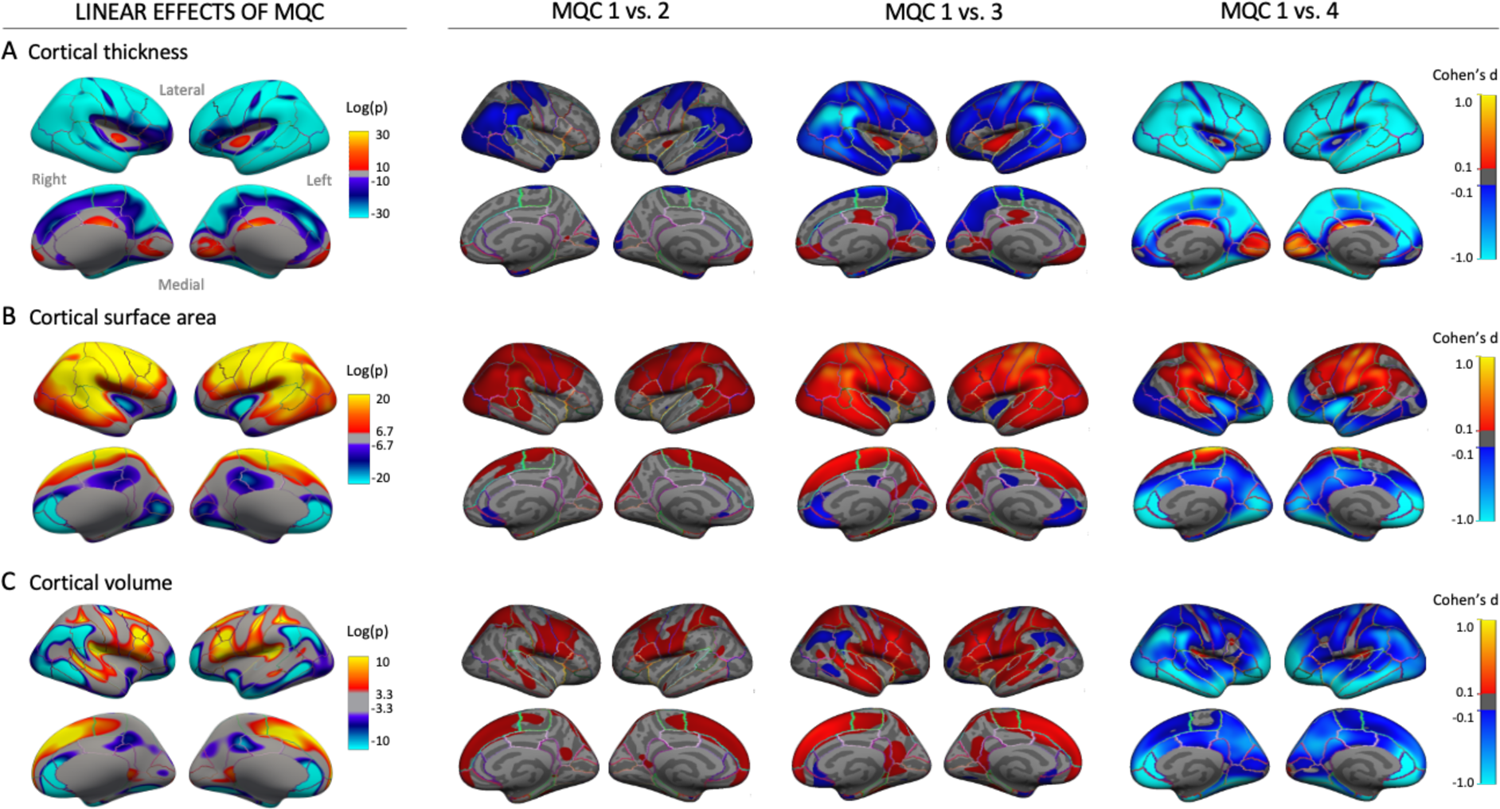
Association between MQC ratings and sMRI indices, n=10,261. Maps at left show linear associations of MQC rating (1 to 4) with cortical thickness (A), surface area (B), and volume (C). Maps at right contrast thickness, surface area, and volume highest quality images (MQC=1) with those assigned to lower quality ratings. Covariates included age, gender, estimated intracranial volume (fixed effects), site, and scanner manufacturer (random effects). Of the initial 10,295 scans with MQC, 34 were excluded due to missing covariates or FreeSurfer processing errors.

All scans had also received automated quality control ratings (pass/fail), available as part of the ABCD NDA. Of the 10,295 scans with MQC ratings, all but 325 were designated as recommended for use; these 325 fell disproportionately within higher MQC groups (comprising 0.4% of MQC=1 scans, 1.4% of MQC=2, 10.6% of MQC=3, and 48.9% of MQC=4) but this designation missed numerous poorer-quality scans. Subsequent analyses including these 10,295 scans were all adjusted for participant age, gender, total intracranial volume, study site, and scanner manufacturer; region-of-interest (ROI)-based analyses were further covaried by family ID to control for effects of participant relatedness.

### Associations between MQC ratings and cortical structure, and comparison with surface hole number (SHN)

Automated measures of cortical thickness, surface area, and volume are commonly used to identify case-control differences or as predictors of dimensional measures (e.g., psychopathology) in psychiatric neuroimaging research ^8^. We next determined the extent to which MQC ratings associate with variance in these measures, as determined by FreeSurfer. MQC ratings associated linearly with reduced thickness across much of the cortical mantle (**Figure 2a**), with increased cortical surface area in lateral/superior and reduced surface area in medial/inferior regions (**Figure 2b**), and heterogeneous effects on cortical volume (**Figure 2c**). Pairwise comparisons of best quality (MQC=1) versus lower quality (MQC=2, 3, and 4) images demonstrated increasingly strong effects on each structural index as QC ratings worsened, with moderate to strong effect sizes noted in numerous cortical regions (see also **Table S3a, b, c** for effects of MQC rating differences in each of the 68 cortical ROI defined by the Desikan-Killiany Atlas). For example, comparison of cortical thickness values between MQC=1 versus MQC=2, 3, and 4 yielded a total of 39, 55, and 61 ROI (of 68), respectively, with statistically significant differences (FDR q <.05). Regions demonstrating stronger effects of poor quality control on thickness included, but also extended beyond, those identified as showing similar effects in a previous, smaller study of adolescent and adult participants (n=1,840) ^5^, in consistent directions (e.g., increased thickness in numerous lateral ROIs, decreased thickness in medial occipital and posterior cingulate cortices). Subcortical volumes also differed significantly based on MQC rating, with higher ratings generally associated with smaller volumes (**Table S4**).

We next compared the performance of an automated QC measure, the surface hole number (SHN), to manual (MQC) ratings. SHN reflects the Euler number, which measures continuity of tessellated images (e.g., those that contain continuous triangular structures, as do FreeSurfer-generated maps of the cortical surface, see **Methods**) based on the sum of the vertices and faces subtracted by the number of faces. Higher SHN have predicted worse manual quality control ratings in previous MRI studies and have been proposed as an automated quality control index for use in high-throughput neuroimaging studies, outperforming other measures (such as signal-to-noise ratio and motion during functional MRI scans conducted during the same scan session) ^5, 16^. We calculated SHN for each available Baseline and Year 2 scan using FreeSurfer 7.0 and have uploaded the data to the NDA (see **Data Availability**). SHN increased in tandem with MQC ratings (rho=0.59; mean SHN differed between all MQC level pairs, p≤1.02E-121), and linear associations of SHN with differences in cortical thickness, surface area, and volume (**Figure 3a,b,c**) closely resembled those of MQC (**Figure 2**). Distribution of SHN values among MQC groups was stable over the temporal sequence of MQC evaluations (**Extended Data Figure 1b**).

**Figure 3.**
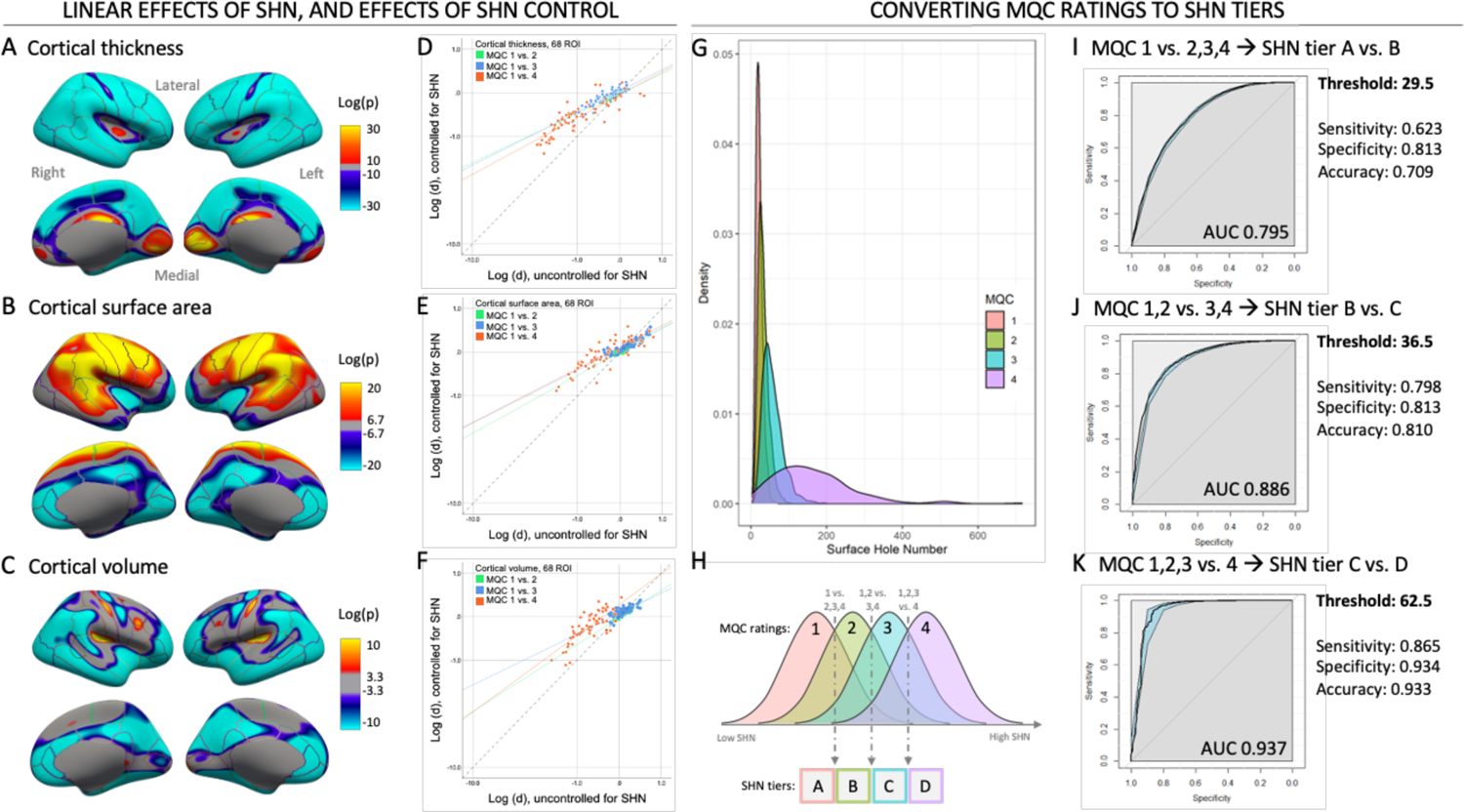
Effects of surface hole number (SHN) on sMRI indices, and derivation of SHN tiers in conjunction with MQC ratings, n=10,261. Linear associations of SHN (non-transformed) with cortical thickness (A), surface area (B), and volume (C) closely resembled those of MQC ratings (compare to Figure 2). Covariates included age, gender, estimated intracranial volume (fixed effects), site, and scanner manufacturer (random effects). Additional adjustment for SHN diminished the effect sizes of pairwise MQC contrasts for thickness (D), surface area (E), and volume (F). Markers represent effect sizes for pairwise MQC contrasts in each of 68 cortical regions-of-interest, and solid lines reflect best-fit across all 68 regions for a given pairwise MQC contrast. Note reduced slopes compared to dashed unity line. (G) Density plot of SHN values, stratified by MQC ratings. Panel (H) illustrates the overall approach for deriving SHN tiers from MQC ratings. The SHN tiers were developed to provide quality control estimates in the absence of manual ratings, and are based on optimized SHN thresholds for parsing higher versus lower manual quality scan groupings. Receiver-operating characteristic (ROC) analyses for various thresholds are shown in panels (I), (J), and (K) along with related specificity, sensitivity, and accuracy indices. For example, with an optimized SHN threshold of 29.5, 81.3% of scans with MQC=2 and higher are eliminated. This threshold was used as a breakpoint for SHN tiers A and B. Blue shaded regions indicate 95% confidence intervals. AUC: area under the curve.

We then examined whether including SHN as an additional covariate mitigated effects of variable scan quality on sMRI indices, as defined by differences in measurements between MQC=1 and MQC=2, 3, and 4 respectively (**Table S3d,e,f; Figure 3d,e,f**). Depending on the specific comparison (MQC=1 vs. 2, 3, or 4), inclusion of SHN reduced the effect size (Cohen’s d) of manual quality control-related differences in cortical thickness by 42 to 59%; reductions in effect size for cortical surface area ranged from 39 to 57%, and for cortical volume from 16 to 62% (**Table S5**). Meaningful effects of SHN correction can also be demonstrated by comparing the number of ROIs that showed statistically significant (FDR, q<.05) effects of MQC ratings before versus after including SHN as a covariate. For example, among 39 ROIs exhibiting differences in cortical thickness between MQC=1 and MQC=2 before covarying for SHN, 17 fell out of significance after covarying for SHN, while 1 ROI became newly significant.

We then used SHN data in concert with MQC ratings to develop and assess the reliability of a tiered, automated sMRI QC rubric to classify the quality of individual scans. This rubic assigned scans to 4 levels akin to the MQC groups, but based exclusively on SHN-based thresholds, so that these ratings could be applied even in the absence of manual QC. **Figure 3g** displays the distribution of SHN among MQC groups. Using receiver operating characteristic (ROC) curve analyses, we derived 3 optimized SHN thresholds to isolate poorer-quality scans (**Figure 3h**). The most conservative threshold eliminated scans with MQC ratings of 2 or higher, based on an SHN cutoff of 29.5 (sensitivity=0.81; **Figure 3i**). The next threshold eliminated scans with MQC ratings of 3 or higher, based on a SHN cutoff of 36.5 (sensitivity=0.81; **Figure 3j**). The most liberal threshold eliminated scans with MQC ratings of 4, based on an SHN cutoff of 62.5 (sensitivity=0.93, **Figure 3k**).

These 3 thresholds defined 4 SHN groups (tiers A-D), that in turn associated with linear effects on sMRI indices (**Extended Data Figure 3**). The linear effects of SHN tiers closely approximated the linear effects of MQC groupings (**Figure 2**), as well as those of continuous SHN values (**Figure 3a,b,c**). Still, MQC and SHN each accounted for distinct variance in scan quality as seen in **Extended Data Figure 4**. In a sensitivity analysis, inclusion of scans with FreeSurfer segmentation errors (n=228) did not substantially alter either the distribution of SHN across MQC ratings or optimal boundaries between SHN tiers in ROC analyses (**Table S6**).

**Figure 4.**
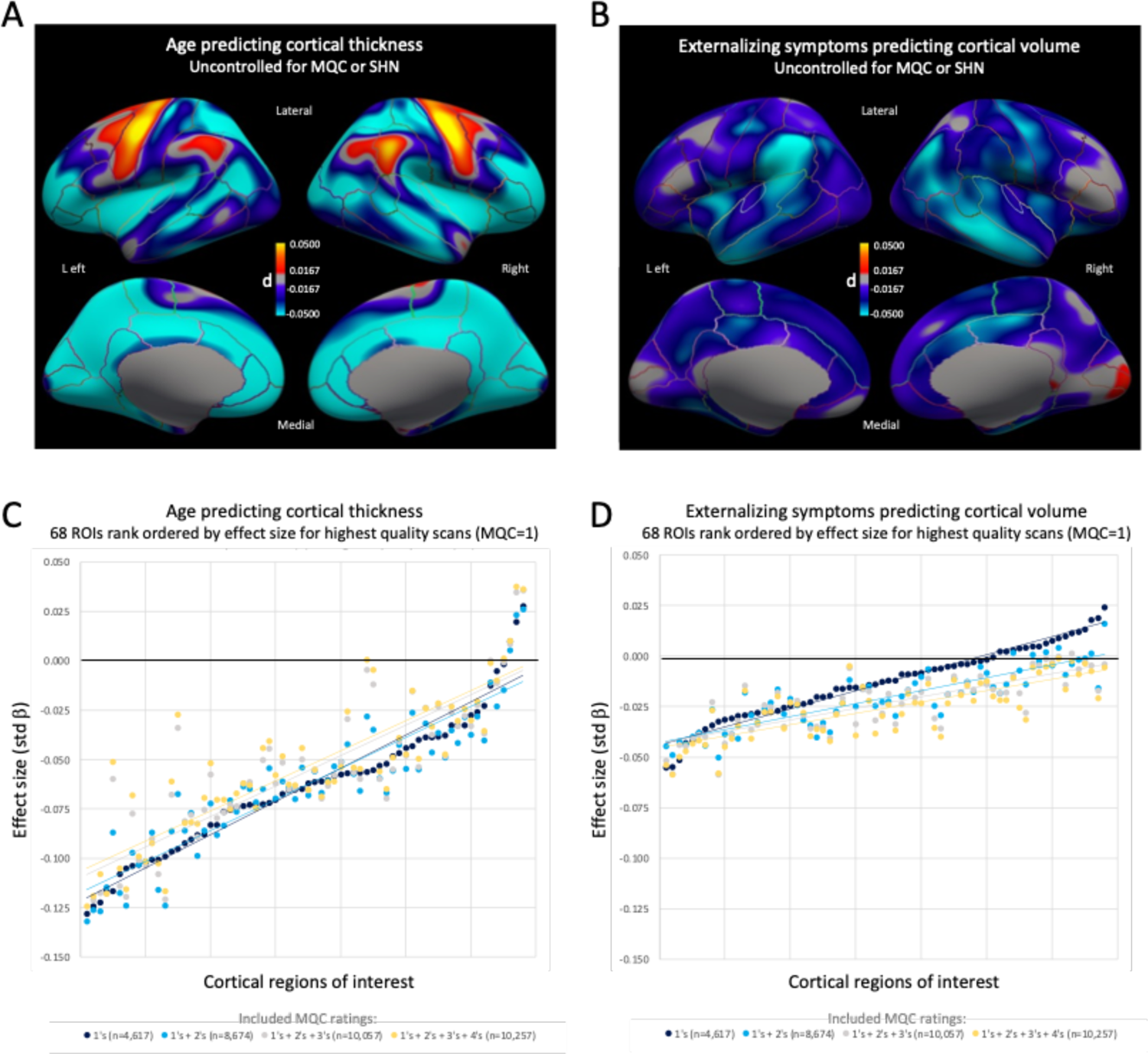
Effects of variable quality control on applied analyses of sMRI data. (A) Association of age with cortical thickness, without adjusting for manual quality control (MQC) rating or surface hole number (SHN). Note the substantially smaller effect size scale compared to Figure 2, which shows effects of quality control variance on sMRI measurements. (B) Association of externalizing symptoms (CBCL externalizing subscale) with cortical volume, without adjusting for MQC or SHN. Note the even smaller effect size compared to effects of age on thickness. (C) Age-thickness effects stratified by region of interest (ROI) and MQC inclusion threshold. Broken lines indicate best-fit lines across all ROIs for each inclusion threshold group. Note tendency toward *diminished* effect size (and increased risk for false negatives) with broader inclusion thresholds. (D) Externalizing symptoms-volume effects stratified by ROI and MQC inclusion threshold. Broken lines indicate best-fit lines for each inclusion threshold group. Note tendency toward *inflated* effect size (and increased risk for false positives) with broader inclusion thresholds. All analyses covaried for age, gender, estimated intracranial volume (fixed effects), site, and scanner manufacturer (random effects); ROI-based analyses also included family ID as a random effect.

### SHN tiers as predictors of MQC in Year 2 follow-up scans

Evaluation of Year 2 scans from ABCD enabled us to test the reliability of SHN tiers derived from Baseline scans. A total of 6,941 minimally processed Year 2 T1 volumes were available through the ABCD Data Archive, after removing those that did not meet inclusion criteria for Baseline analysis; see **Extended Data Figure 5**. Following preprocessing in FreeSurfer 7.0 and extraction of SHN, 1,000 sMRI volumes were semi-randomly selected such that they included (1) a range of scan quality, operationalized by ensuring a mix of tiers A, B, C, and D; and (2) a distribution of magnet types (Siemens, Philips, GE) that was equivalent to the analyzed Baseline sample. Of note, Year 2 scans showed better overall quality than Baseline scans, with 83.9% falling into SHN tier A (**Extended Data Figure 6a**; compare to **Figure 3g**, where 57.3% of Baseline scans fell into tier A). Group characteristics of SHN tiers A to D in the Year 2 sample are described in **Table S7**. One scan was discarded due to presence of a large cyst. Only 168 Year 2 scans fell within SHN tier D, all of which were used for the analysis. The selected Year 2 scans underwent MQC ratings by 2 trained and calibrated research coordinators (500 scans randomly disbursed to each of K.A.K. and E.L; see **Methods**), using the same method as Baseline MQC ratings.

**Figure 5.**
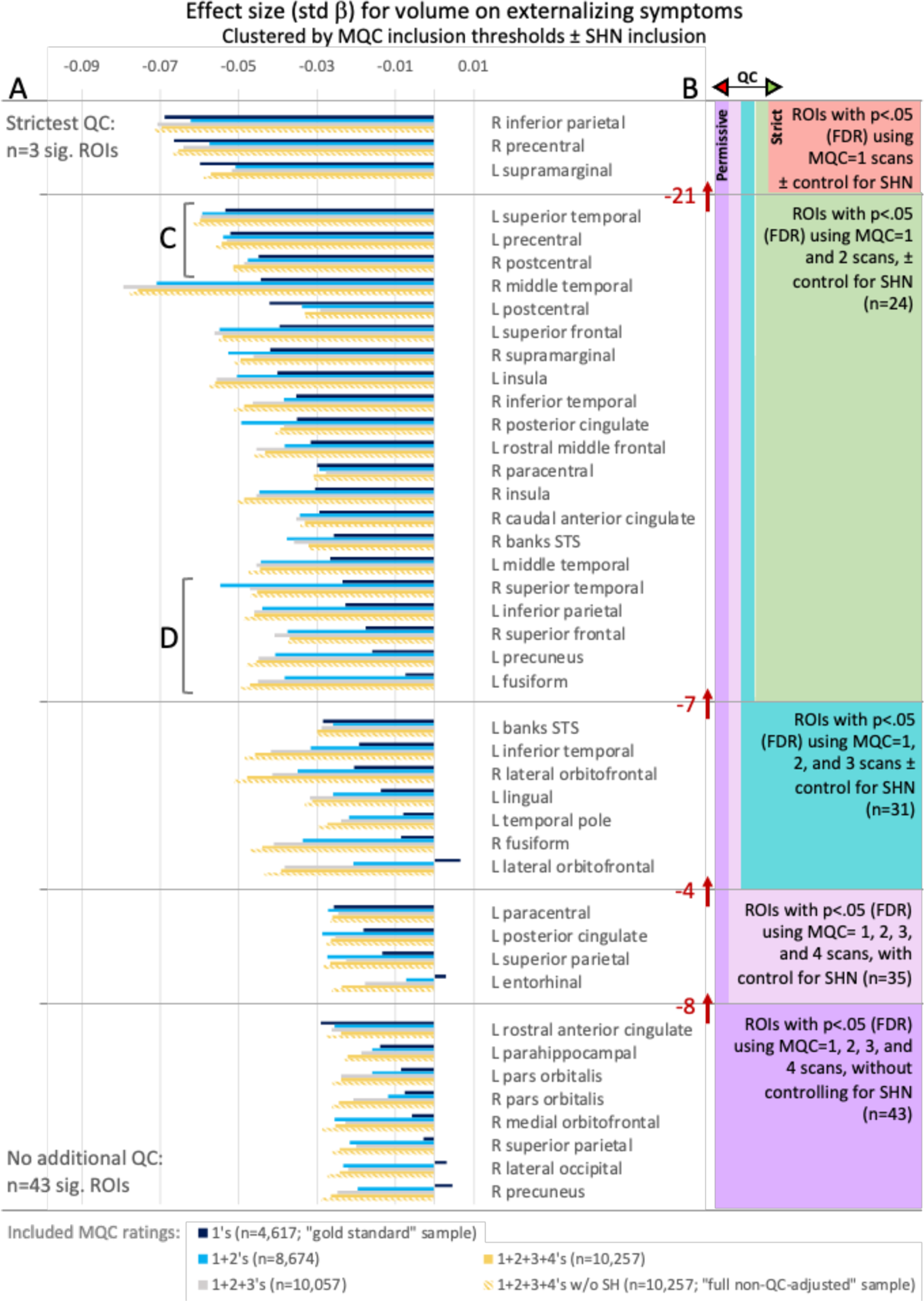
Effects of increasingly stringent quality control on effect size and statistical significance of externalizing symptoms-volume findings. (A) At left, bars indicate effect sizes for the relationships between externalizing symptoms and cortical volume for each ROI, stratified by the stringency of quality control of included scans (see legend). At one extreme, dark blue bars indicate effect sizes generated by using the most conservative approach, i.e., only MQC=1 scans were included in the analysis, which also corrected for SHN (“gold standard” sample, n=4,617). At the other extreme, thatched yellow bars indicate effect sizes generated by using the most permissive approach, i.e., scans with all MQC levels were included in the analysis, and no SHN correction was applied (“full non-QC-adjusted” sample, n=10,257). Note that for most regions, more permissive quality control was associated with inflated effect sizes. (B) As seen at right, ROIs were grouped based on whether they continued to show statistically significant (q<.05, FDR) relationships between externalizing symptoms and cortical volumes as lower quality scans were iteratively removed. Red numbers and arrows indicate the number of ROIs that dropped out of significance with each level of tightened QC. The purple group contains 43 regions that were significant after using the most permissive quality control (no removed scans). In contrast, the red group (gold standard) contains only 3 regions that were significant after using the most conservative quality control (MQC=2, 3, and 4 removed). Note that when including the next best of scans (MQC=2), several regions that become significant in this larger sample (n=8,674), e.g., those near (C), did not show inflated effect sizes when lower quality scans were included, and thus appeared robust to poor scan quality. That these regions are significant when MQC 1+2 scans are included in the analysis – but *not* significant when only MQC=1 scans are included (n=4,617) – indicates that using only the highest quality scans potentially results in false negatives (type II error) due to lack of statistical power. However, other regions that were significant in the MQC 1+2 group, e.g., those near (D), showed substantial effect size inflations when scans rated as MQC=2 or higher were included. For these regions, statistically significant findings likely reflected false positives (type I error) – even when all included scans were of relatively good quality. All analyses covaried for age, gender, estimated intracranial volume (fixed effects), and site, scanner manufacturer, and family ID (random effects).

**Figure 6.**
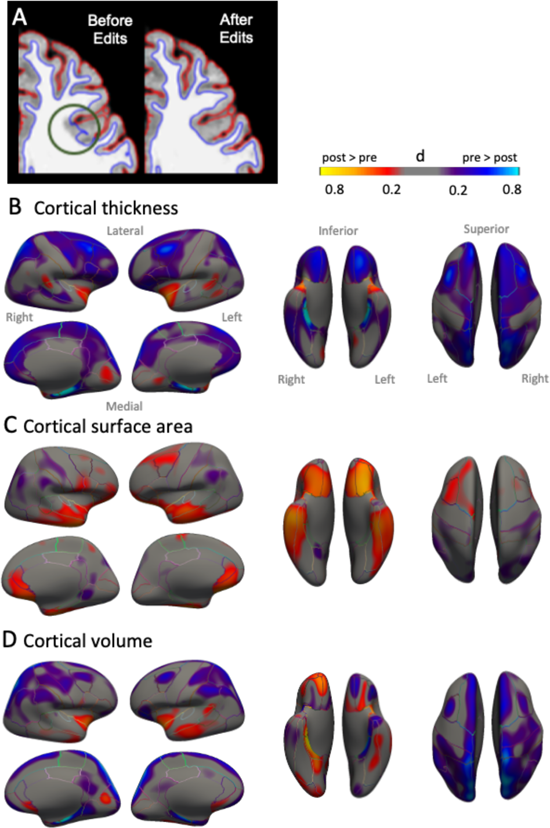
Effects of manual edits on sMRI indices (n=180). Manual edits (e.g., A, which corrects a gray-white matter boundary segmentation error) were conducted on 150 scans with MQC=1 and 30 scans with MQC=2. Maps reflect effect sizes (Cohen’s d) of pre-to-post edit changes in (B) cortical thickness, (C) cortical surface area, and (D) cortical volume.

**Table S8** describes the performance of SHN tiers in predicting MQC ratings for the 999 Year 2 scans. The SHN tiers effectively filtered out scans with higher MQC ratings, with sensitivity ranging from 0.87 (for differentiating scans rated 2 and higher from those rated 1) to 1.00 (for differentiating scans rated 4 from those rated lower). **Extended Data Figure 6b** shows the distribution of MQC ratings within each SHN tier. **Extended Data Figure 7** indicates the effect of SHN tiers on sMRI indices across all 6,941 Year 2 scans (most of which had not received MQC ratings); comparison to **Extended Data Figure 3** affirms that SHN tiers reproducibly tracked variance in scan quality, especially in regard to cortical thickness and surface area.

### Scan quality and risk for error in applied sMRI analyses

sMRI measures are frequently explored for associations with clinical and developmental data. The ABCD Study provides an unprecedented opportunity in this regard, with multiple imaging and clinical measurements obtained within the same youth participants over a 10-year period. However, given the tendency of poorer quality images to bias sMRI measurements among youth, we next examined the extent to which unaccounted variance in scan quality might affect associations between MRI and clinical indices.

As a positive control, we first considered a well-established relationship between age and cortical thickness. Most of the cortex is known to thin linearly during adolescence, as seen in smaller but well quality-controlled samples ^24^. As points of reference, we compared age-thickness effects in the SHN-corrected, MQC=1 sample (n=4,617, “ground truth”) to those in the full, non-corrected sample (n=10,257, “full non-QC-adjusted sample”). Significant age-thickness relationships were readily observed, even cross-sectionally between ages 9.0 and 10.9, within the full non-QC-adjusted sample (**Figure 4a**) – although note the considerably smaller effect size of age on thickness compared to that of quality control ratings (**Figure 2a**). Despite these smaller effects, age-thickness effects were sufficiently robust to be detected within the smaller ground truth sample: among 68 cortical ROIs, significant (FDR q<.05) negative associations were present in 59 regions, regardless of SHN adjustment (**Table S9**). Notably, though, several of these ROI did *not* show significant age-thickness differences in the (larger) full unadjusted sample – but then *regained* significance in the full sample after SHN adjustment. As such, inclusion of SHN mitigated Type II error (i.e., false negatives) that would have otherwise occurred in the full non-QC-adjusted sample, albeit for only a small number of regions.

The risk of Type II error arising from non-quality-corrected images can also be appreciated in **Figure 4b**, which plots effect sizes for the age-thickness relationship across all 68 ROIs. To facilitate comparisons across MQC levels, ROIs were rank-ordered (left-to-right) by effect size among the 1-rated scans. Effect sizes generally diminished as poorer quality images were iteratively included (2s, then 3s, then 4s) in the analysis. These results echo a prior, smaller analysis (n=1,598, mean age=15.0), wherein poorer quality scans associated with blunted effects of age on cortical thickness ^5^.

Next, we considered a more exploratory relationship between dimensional psychopathology and cortical volume. Several groups have reported inverse associations between CBCL scales and cortical volume, including using ABCD data ^25, 26^. In a recent study focused on genetic and neurodevelopmental underpinnings of psychopathology in ABCD ^27^, among the broadband CBCL scales (total, internalizing, externalizing) we identified externalizing symptoms (CBCLext) as most strongly related to cortical volumes at Baseline, after taking into account both MQC ratings and SHN.

In the full, non-QC-adjusted sample, CBCLext scores showed a diffuse, inverse relationship with volume across the cortical mantle (**Figure 4c**), although effect sizes were smaller than for the age-thickness relationship (**Figure 4a**). Within this larger sample, 43 ROIs demonstrated significant (FDR q <.05) relationships between CBCLext and volume (**Figure 4d, Table S10**). However, stark differences emerged in comparison to the ground truth (MQC=1) sample, wherein only 3 regions demonstrated significant CBCLext-volume relationships. Stepwise analyses that gradually increased the stringency of QC suggested that this drop in the number of significant ROIs reflected an interplay of QC and power considerations (**Figure 5**). While effect size should not depend on sample size, unlike in the age-thickness analysis, inclusion of lower-quality images resulted in substantial inflation of CBCLext-volume effects, and accordingly Type I errors (i.e., false positives). Numerous ROIs showed statistically significant CBCLext-volume relationships only when MQC=3 and 4 scans were included in the analysis – and, even after correction with SHN, these regions demonstrated inflated effect sizes due to inclusion of lower quality scans. Further, regions with smaller effect sizes in the ground truth sample were more likely to show inflated effect sizes – and, hence Type I error – in the full, non-adjusted sample (**Figure 4d**). This result was counterintuitive, given that large sample size is often invoked to *reduce* risk of *Type II* error, through improved power to detect small but true effects.

Further complexity emerged among the 21 ROIs that showed significant CBCLext-volume relationships after including only MQC ≤ 2 images. For some regions, such as left superior temporal, left precentral, and bilateral postcentral, effect size remained relatively stable as inclusion thresholds loosened (**Figure 5c**). This pattern suggests that effect sizes were not inflated by artifact, and that failure to reach significance when using only MQC=1 scans reflected a lack of statistical power (i.e., Type II error) – even with a sample size of >4,500. For others, such as right middle temporal, bilateral insula, and bilateral superior frontal, effect size increased substantially as the MQC inclusion threshold was relaxed to 2 (or higher), likely reflecting artifact. For these regions, inclusion of even relatively good quality (but not the highest quality) images appeared to result in Type I error (**Figure 5d**).

### Effects of manual edits on sMRI indices

Image reconstruction errors can influence sMRI measurements and can be exacerbated by head motion and other artifacts ^3, 4^. These errors include skull strip errors, segmentation errors, intensity normalization errors, pial surface misplacement, and topological defects. Within FreeSurfer these errors can be corrected through manual editing of voxels in brain and white matter masks, watershed thresholds, and the addition of control points ^15, 16^. Here, we examined effects of manual edits on sMRI indices among scans with relatively higher image quality, to assess whether this intervention might safely be reserved for those with MQC >2.

A total of 150 Baseline scans with MQC=1 and n=30 Baseline scans with MQC=2 were randomly selected for manual edits by a trained coordinator (see **Methods**). Direction and effect sizes of pre-to-post edit changes across the cortical mantle are displayed in **Figure 6** (MQC=1 and 2 combined, n=180) and **Extended Figure 8a** and **b** (MQC=1 and 2 separately), while ROI-level changes across the entire sample of 180 are described in **Table S11a,b,c**. Effects of manual edits were most pronounced for cortical thickness and volume, both of which tended to decrease. These changes reached statistical significance (FDR q <.05) for cortical thickness in 40 regions (Cohen’s d range 0.16 to 0.92), and for cortical volume in 28 regions (d range 0.18 to 0.73). Numerous regions with signal across all scans (MQC=1 and 2) demonstrated stronger effects of editing on cortical thickness and volume in MQC=2 scans than MQC=1 scans (e.g., bilateral parahippocampal, caudal middle frontal, and superior parietal cortices). Further, cortical volume maps revealed a strong effect of edits in the area of the superior sagittal sinus, particularly impacting superior parietal cortex (**Extended Figure 8c**). In an applied analysis, we then examined the degree to which cortical edits affected effect size for the relationship between cortical thickness and age. Across all 68 cortical ROIs, effect size slightly strengthened (became more negative) for post-edited images compared to pre-edited images (t=2.31, p=0.024, d=−0.10).

To put these findings in context along with MQC rating effects on sMRI indices, **Extended Figure 9** maps all ROIs that showed significant (FDR, q <.05) effects of MQC, surface edits, or both, as well as their direction, among Baseline scans with MQC=1 or 2. Note that even when constrained to the best two scan quality groups, there are diffuse effects of scan quality differences across the cortex, for each of the sMRI indices; and that biases related to poorer overall quality control and to subtle topological defects can induce opposing effects on sMRI measurements.

Finally, to assess reproducibility and developmental specificity of cortical edit effects, we compared ABCD results to that of a second, non-overlapping MRI cohort of 292 youths, age 8 to 18, who received MRI scans that were assessed by radiology reports as free of pathology at Massachusetts General Hospital (MGH; **Table S12a, b, c**). This sample was previously described in an analysis relating prenatal folic acid exposure to cortical development^20^. This sample differed from ABCD by its inclusion of (1) clinical rather than research participants, (2) *all* editable images (not just those of relatively high overall quality), (3) a mix scanner field strengths (1.5 and 3T) as well as manufacturers, and (4) a broader age range. Despite these differences, of the 40 regions demonstrating significant effects of manual edits on thickness in ABCD, 18 again showed nominally significant (15 showed FDR-significant) effects of edits in the same direction within the MGH MRI cohort (Cohen’s d range 0.12 to 0.98). Notably, across these 18 regions, differences in pre-to-post edit mean thickness were greater at age 8-10 compared to other age groups (11-12, 13-14, and 15-17; omnibus F=8.49, p=0.0001, post hoc comparisons p’s≤.0002; **Extended Figure 10a**). Similarly, the standard error of pre-to-post thickness changes across individuals was also greatest at age 8-10 (omnibus F=64.53, p=2.25E-17, post hoc comparisons of age 8-10 vs. other groups, p’s≤6.53E-10). Finally, the effect of edits on the relationship between age and cortical thickness differed among age groups (F=21.54, p=3.88E-12); specifically, the effect of edits on the age-thickness relationship was stronger at age 8-10 (d=-1.18) than for any other age group (p’s≤7.73E-09; **Extended Figure 10b**). These results indicate that manual edits result in replicable, diffuse changes in cortical thickness in early adolescence that can influence effect sizes in clinical-MRI associations, but also suggest that effects of edits become less pronounced later in adolescence.

## Discussion

### Implications for brain-wide association studies in youth

These present findings identify nuances related to scan quality in large pediatric brain MRI cohorts that are pervasive and complex, and that likely require multi-pronged intervention to avoid error in MRI-based analyses. Leveraging one of the largest collections of uniformly collected sMRI data from children and adolescents, we used manual quality control (MQC) to separate high quality scans and contrast them to those with various degrees of observable artifact. While inclusion of lower-quality scans diminished variance in estimates of widely used sMRI metrics, such as cortical thickness and surface area, they also introduced substantial bias. These effects were partially mitigated by inclusion of surface hole number (SHN), an automated measure of topological complexity that accounted for quality-related variance in sMRI measures akin to MQC. However, inclusion of SHN failed to safeguard against most Type I and II errors when poorer quality scans were included in applied analyses that associated sMRI measures with clinical data. Further, even among the highest quality scans, manual editing associated with significant changes in cortical thickness and surface area -- changes that in some regions were oppositely signed to those observed when controlling for SHN or MQC, and that replicated in a non-overlapping clinical cohort. As a whole, these results challenge assumptions that large sample size alone improves sensitivity to detect valid brain-behavior relationships, or mitigates the effects of variable image quality on error risk.

Implications of these findings extend not only to studies that map trajectories of healthy and aberrant brain development, but also to applied analyses that relate structural indices to clinical measures. Comparison of effect sizes for sMRI-clinical relationships (**Figure 5**) to those of bias related to poor scan quality (d=0.14-2.84) or manual edits (d=0.15-0.92) – which are generally higher by an order of magnitude – demonstrates the susceptibility of these relationships to artifact. Recent analyses illustrate the need to include thousands of individuals in brain-wide association studies ^28, 29^, reflecting the small effect sizes intrinsic to these relationships. Here, inclusion of the best quality (MQC=1, n=4,617) scans was inadequate to detect relationships between cortical volume and externalizing psychopathology in several regions, effects that became statistically significant when scans of marginally lower quality (MQC=2, n=4,057) were included. However, further inclusion of even lower quality (MQC=3 and 4, n=1,585) scans resulted in statistically significant but errant associations between volume and externalizing psychopathology in dozens of brain regions, as demonstrated by markedly inflated effect sizes compared to analyses that included exclusively higher quality scans. These results indicate complex trade-offs between sample size and scan quality that warrant careful consideration in large MRI studies, especially in the setting of small effect sizes.

### Quality control in the era of “big data” sMRI studies

Large and diverse study samples offer clear advantages such as statistical power and improved generalizability, and in the case of psychology and neuropsychiatry research, such designs help to mitigate well-described problems of publication bias and reproducibility failures ^28, 30^. However, several pitfalls within “big data” science have also been described, including inadequate control for multiple comparisons, sampling bias, measurement error, and discrepancies between statistical and clinical significance. These issues have hampered other areas of clinical-translational research, such as electronic health record, epidemiology, and health services studies ^31^. With regard to brain imaging research, a recent study ^32^ used theoretical data to model trade-offs of increasing sample size well into the thousands, demonstrating the risk of latent bias to outweigh the benefit of reduced variance. This concern bears out in the present real-world analysis, which cautions against equating data quantity and quality in youth sMRI studies ^33^. The findings also have important implications for large-scale MRI studies of other populations where head motion occurs more frequently, including those with psychiatric and neurological disorders ^34^, and those at the extremes of age ^35, 36^.

Beyond best practices to minimize participant motion^23^, the present findings suggest that relatively labor-intensive approaches – visual QC and manual editing – conducted in concert with automated measures such as SHN – provide the best protection against errant sMRI findings in youth cohorts. However, manual edits pose their own challenges with regard to feasibility in studies with tens of thousands of participants, as editing each poorer quality scan can require hours of personnel time. The present analyses offer several QC alternatives that may be weighed in the context of available resources, the nature of particular findings, and the characteristics of the study population. For example, the SHN benchmarks identified and validated in this report provide an alternative QC approach that is imperfect but less time-consuming. Investigators who find associations of sMRI indices with behavioral, genetic, or environmental factors described in ABCD and other neurodevelopmental cohorts may wish to consider both the effect sizes and anatomical distribution of these associations in deciding which QC approach is appropriately conservative. The present analyses of older adolescents (ABCD Year 2, MGH) are encouraging and suggest that less intervention may be needed with advancing participant age. Further, as automated methods continue to gain sophistication ^37, 38^ they may continue to improve the efficiency of QC and further strengthen causal inference in neurodevelopmental MRI research.

## Methods

### Sample from ABCD

The ABCD Study has collected data from 11,875 children from 22 sites across the United States. Primary analyses used baseline data from children aged 9-10 years old. Institutional Review Board (IRB) approval for the ABCD study is described in Auchter et al. ^39^ All parents provided written informed consent and all youth provided assent. We excluded subjects whose baseline MRI scans were flagged for clinical consultation (N=451), and those without available T1 data (N=160) from all analyses.

### MRI acquisition

All MRI images were obtained using harmonized parameters with 3T MRI scanners manufactured by Siemens, Philips, or GE. We used T1 weighted images (256×256 matrix, slices=176-225, TR=6.31-2500, TE=2-2.9, 1×1×1 mm resolution) for our analysis. Images acquired from Siemens and GE scanners included real-time motion detection using volumetric navigators that automatically triggered re-scans ^11, 12^. Additional details of MRI sequences are described elsewhere ^23^.

### Image processing

Minimally processed baseline T1 images from 11,264 participants, and year 2 follow-up T 1 images from 6,941 of these participants, were downloaded from the ABCD Data Archive (release 4.0). Scans underwent N4 field bias correction to correct low frequency intensity non-uniformities or field bias ^40^. Subsequently whole brain processing and analyses were conducted using FreeSurfer version 7.1 (http://surfer.nmr.mgh.harvard.edu/). One baseline scan failed Freesurfer processing. Using automated segmentation (Desikan-Killiany atlas), cortical thickness, surface area, and volume of 68 regions of interest (ROI) were extracted, as were 20 subcortical volumes.

### Manual quality control (MQC)

A single, trained rater (S.E.) conducted visual assessment of all processed Baseline scans. The rater was blinded to any potential identifying, clinical, or demographic information regarding participants. This rater had been trained by the PI (J.L.R.) and a clinical research coordinator (K.F.D.) who had experience conducting manual edits of >300 MRI scans acquired from children and adolescents aged 8 to 18 ^20^. The system of rating was developed by using a randomly selected set of 200 baseline T1 scans. The final manual quality control (MQC) ratings scheme, developed in consensus with S.E., K.F.D., and J.L.R., included 5 categories: A rating of “1” was given to scans of minimal artifact, only needing about ½ hour to complete edits. A rating of “2” was given to scans with moderate artifact, requiring 1-2 hours for manual edits. A rating of “3” was given to scans with substantial artifact, requiring several hours of edits. A rating of “4” was given to scans with severe artifact, such as motion artifact, and would not be possible to fix with manual edits. Lastly, a rating of “5” was given to scans with a processing defect which resulted in segmentation errors and apparent loss of tissue. Scans that included cysts that were greater than 1 cm^3^ were not rated and excluded from subsequent analysis. The order in which scans were evaluated for MQC was semi-random. Scans originating from N=5,105 participants of European ancestry were prioritized and randomly sequenced to facilitate a genomic analysis^27^. However, this initial group also contained 373 randomly interspersed scans from randomly selected non-European participants. Following assessment of this initial set of 5,105 scans, the remainder were evaluated in random order. Of the evaluated scans, 368 were coded within the ABCD NIMH Data Archive as “inclusion not recommended” based on an automated overall QC measure in the FreeSurfer preprocessing stream and/or corrupted raw data at the time of scan acquisition (imgincl_t1w_include=0); the remainder received the “inclusion recommended” code.

### Characterizing apparent tissue loss due to segmentation errors

MQC=5 scans (N=228) were re-rated as MQC from “1” through “4” to assess the quality of the remaining volume that was unaffected by segmentation errors. Ratings were performed by the same trained rater who assigned ratings to all baseline scans. The sagittal, coronal, and axial extents of the drop out region were measured in Freeview (https://surfer.nmr.mgh.harvard.edu/fswiki/FreeviewGuide). Approximate volumes of segmentation error-related tissue loss were calculated assuming an ellipsoid shape and measured x, y, and z dimensions. For purposes of displaying location and overlap of drop-out across scans, rectangular cuboids were constructed in MarsBar in SPM 12 (http://www.fil.ion.ucl.ac.uk/spm/) using measured dimensions and coordinates. Rectangular cuboids were combined across subjects and were thresholded by a whole brain mask in MarsBar. Areas of dropout were thresholded at n>10 subjects with dropout in that region and drop-out was displayed on an exemplar structural image in xjView (https://www.alivelearn.net/xjview).

### Surface hole number

We used surface hole number (SHN) as an automated quality control measure extracted from FreeSurfer aparc tabulated data. SHN is a topological measurement referring to geometrical holes (imperfections) in the tessellated brain surface. SHN is related to the Euler number by the formula, Euler number = 2 - 2 × SHN. Previous, smaller studies have suggested that SHN can serve as a proxy for overall T1 scan quality ^5, 16^. Here, SHN from baseline scans were used to determine optimal proxies for MQC, through creating of 4 tiers (A, B, C, D) that approximated the 4 levels of MQC ratings (1, 2, 3, 4).

### Psychopathology measurement

We used the parent-reported Child Behavior Checklist (CBCL) as a measure of dimensional psychopathology. The CBCL is a frequently used scale comprising eight subscales (anxious/depressed, withdrawn/depressed, somatic, social, thought, attention, rulebreaking, and aggressive symptoms) that can be summarized by total, internalizing, and externalizing scores. Raw scores are converted to t-scores which are normed for age and gender ^41^.

### Year 2 T1 replication

We examined all available Year 2 T1 weighted images to assess the reliability of SHN tiers derived from Baseline scans. The most recent ABCD data release (4.0) contains Year 2 scans from 7,829 participants. Using the same method as for Baseline scans, we used FreeSurfer to process images from 6,941 individuals whose baseline image passed the inclusion criteria and received MQC ratings of 1-5. SHN were calculated by FreeSurfer for each of these scans. In addition, 1,000 Year 2 scans were semi-randomly selected for MQC ratings, such that they contained (1) a range of scan quality, operationalized by selecting for an approximately equivalent number of scans that fell into tiers A, B, C, and D; and (2) a distribution of magnet types (Siemens, Philips, GE) that was equivalent to the analyzed Baseline sample. These scans were then rated for MQC in random sequence by two raters (E.L, K.A.K.) who had previously been trained by the rater of all Baseline scans (S.E), such that the three raters achieved an intraclass coefficient of >0.75 (two-way mixed effects model for absolute agreement) across a training set of 1,000 Baseline scans.

### Manual cortical edits of ABCD scans

A subset of the rated Baseline scans was randomly selected for manual editing (N=150 with MQC=1, N=30 with MQC=2). Each structural scan was loaded into Freeview version 7.1.1 with the following volumes: brainmask, wm, brain.finalsurfs.manedit, T1, and the following surfaces: rh.pial, rh.white, lh.pial, lh.white. The scans were primarily displayed in the coronal view, although sagittal and horizontal views were used as needed. Criteria for editing were primarily based off overestimation and underestimation of the pial and white matter boundaries. Edits to the white matter boundary were made directly on the wm volume using control points and the erasing tool. Edits to the pial surface were made on the brainmask volume. Errors between the pial surface and cerebellum were corrected using the brain.finalsurfs.manedit volume. Edits were considered to be complete when, after post-edit re-processing in FreeSurfer, there appeared only minimal errors remaining, meaning the generated pial and white matter boundaries more closely matched the actual boundaries on the T1 image.

### Manual edits of Massachusetts General Hospital (MGH) scans

The MGH sample was included as a replication set for effects of manual editing on cortical MRI indices and to assess whether such effects change later in adolescence. Study sample, scanner characteristics, and editing methods were previously described by Eryilmaz and colleagues ^20^. Study procedures were approved by Partners Human Research Committee, which granted a waiver of informed consent, since this retrospective study of the medical record involved only deidentified data. Briefly, clinical brain MRI scans from 292 individuals aged 8 to 17, conducted at MGH between 2005 and 2015, were selected based on date of birth, adequate scan quality on visual inspection (i.e., artifacts could reasonably be addressed with manual edits), and absence of pathology as indicated on radiology reports. Scans were edited by a trained research coordinator (KFD) as described above. Pre-to-post edit changes in cortical thickness, volume, and surface area were measured across 68 ROIs using FreeSurfer 5.0.

### Statistical analysis

#### Stability of MQC ratings over time

MQC ratings of baseline scans that did not show signal dropout or cysts were divided into 10 equally sized time groups, reflecting the sequence in which scans were evaluated. Initial analyses were conducted to assess whether factors known to affect scan quality, including age, gender, scanner manufacturer, and psychopathology (CBCL) differed over time, using time period as either a categorical or continuous variable. Then, ANOVA was used to assess significant linear or quadratic changes in mean MQC rating across time groups, controlling for variation in these other factors and in their interactions with time and time-squared.

#### Surface-based sMRI analyses

Surface maps for group-based and within-subject analyses were generated using Freesurfer 7.0. Images from each subject were smoothed by 22mm full width-half maximum. For between-group analyses we fit general linear models with following covariates: age, gender, estimated intracranial volume, study site, and scanner. Continuous predictor variables were z-transformed prior to analysis. Models assessed both linear effects of MQC ratings (i.e., 1 to 4) as well as pairwise contrasts (1 vs. 2, 1 vs. 3, 1 vs. 4) on cortical thickness, surface area, and volume. Sensitivity analyses assessed linear effects of SHN on these indices, as well as effects of MQC after controlling for SHN and vice versa. Results were visualized using uncorrected significance maps (log p-value) and effect size maps (Cohen’s d) as appropriate.

#### ROI-based sMRI analyses

Following extraction of ROI-based data from Freesurfer, analyses involving cortical thickness, cortical surface area, and cortical and subcortical volumes were conducted with R version 4.1.2 (https://www.R-project.org/). Mixed-effects linear regression was run with “lme4” package (https://github.com/lme4/lme4/), unless specifically mentioned. The covariates included in the analysis were age, gender, estimated intracranial volume (fixed effect), site, scanner, and family ID (random effects), the latter accounting for inclusion of sibling groups. Analyses were corrected for multiple comparisons using FDR (q<.05), based on the number of included ROIs.

#### SHN tiers

We conducted receiver operating characteristic (ROC) analyses to evaluate the sensitivity of SHN to detect poorer quality scans. Analyses were conducted in R using the “pROC” package. Using Baseline scan data, we contrasted SHN for three breakpoints: MQC=1 vs. 2, 3, and 4; MQC=1 and 2 vs. 3 and 4; and MQC=1, 2, and 3 vs. 4. We used the Youden Index to select an optimal threshold to discriminate higher versus lower quality scans for each of the three breakpoints. These three thresholds were used to define SHN tiers A, B, C, and D, respectively – such that scans in the A tier best represented MQC=1, those in the B tier best represented MQC=2, etc. As a sensitivity analysis, we also included MQC and SHN values for scans with segmentation-related tissue loss into the analysis, and examined whether thresholds were altered by inclusion of these scans. Then, to test reliability, we grouped all available Year 2 scans according to SHN tiers, and conducted MQC on 1,000 of these scans (described above).

Sensitivity, specificity, and accuracy of the SHN tiers to distinguish MQC levels were assessed. These metrics could then be compared to those from the Baseline analysis, as well as to those from a new set of ROC analyses that determined optimal thresholds for SHN tiers in the 1,000 Year 2 scans.

#### Applied analyses relating quality control to MRI-clinical associations

Linear mixed models examined associations between cortical thickness and age, and between cortical thickness and externalizing psychopathology, conditioned on the degree to which lower-quality scans were included in the analyses (e.g., inclusion of MQC=1 only, versus MQC 1 and 2; 1, 2, and 3; and 1, 2, 3, and 4). Overall surface-based and ROI methods were similar to those described above, but now using age or CBCL externalizing score rather than MQC as the predictors of interest. Sensitivity analyses examined effects of including SHN as an additional predictor in the models.

#### Effects of manual edits on sMRI indices

For Baseline ABCD scans, within-subject analyses that contrasted cortical thickness, surface area, and volume before vs. after manual edits were conducted using general linear models in Freesurfer (for surface maps of effect size) or paired t-tests in R (for ROI analyses). These analyses were conducted without covariates, following upon sensitivity analyses that demonstrated no significant effects of age, gender, scanner, or CBCL externalizing symptoms on pre-to-post edit changes in sMRI measures. ROI analyses were corrected for multiple comparisons using FDR (q<.05), based on the number of included ROIs. Analyses of MGH scans focused on cortical regions that replicated significant effects of manual edits on cortical thickness that were seen in the ABCD cohort. Potential changes in magnitude and variance of pre-to-post edit changes across these regions were assessed as a function of age group (8-10, 11-12, 13-14, 15-17 years) using ANOVA.

#### Data availability

Data from all ABCD-related analyses were downloaded from the NIMH Data Archive (NDA), version 4.0. Derived variables, including MQC ratings and SHN, as well as region-of-interest level data for cortical thickness, surface area, and volume processed in FreeSurfer 7.0, have been uploaded to the NDA (Study ID #1944, doi 10.15154/1528507). Data from MGH analyses contain sensitive patient information that was obtained following a waiver of informed consent, and as such has not been uploaded to a publicly available repository. Please contact the corresponding author for additional information.

---

**Extended Data Figure 1.**
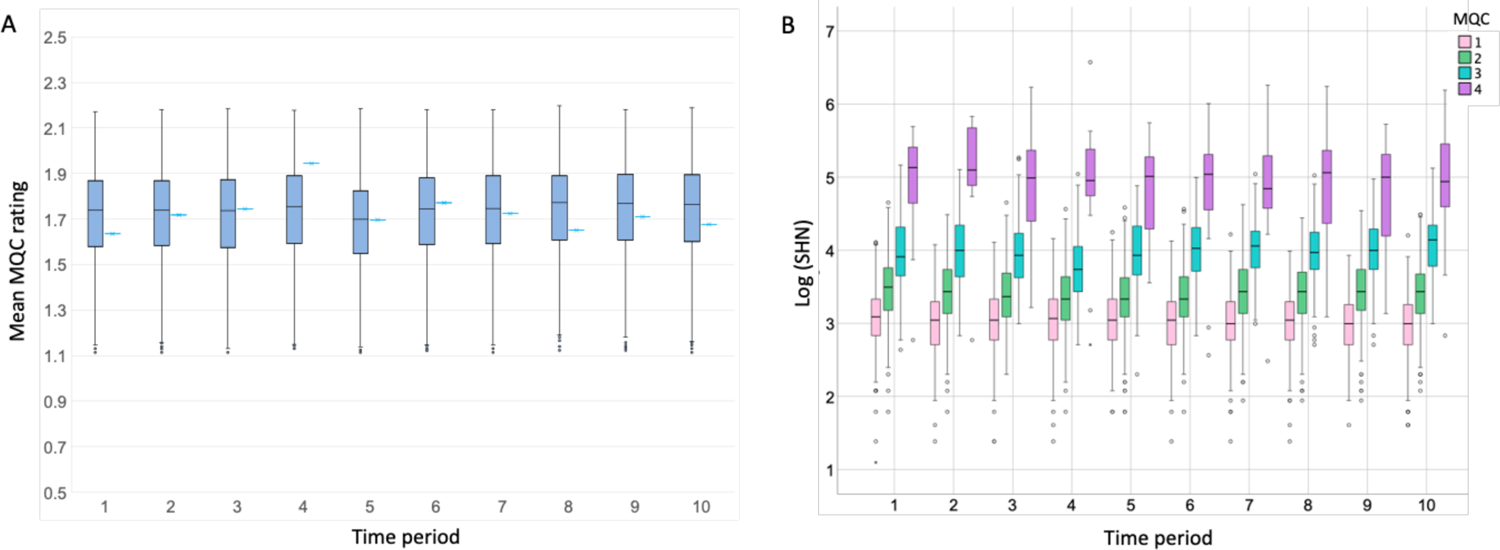
Stability of manual quality control (MQC) ratings over time (n=10,295). Scans were assigned to deciles based on the sequence in which they received MQC ratings by a single trained rater. (A) Box and whisker plots show distribution of MQC ratings for each time period, after adjusting for age, gender, scanner manufacturer, and externalizing psychopathology. Adjacent marks show unadjusted mean ratings for the same period. (B) Box and whisker plots show distribution of the log of surface hole numbers (SHN), stratified by decile and MQC rating.

**Extended Data Figure 2.**
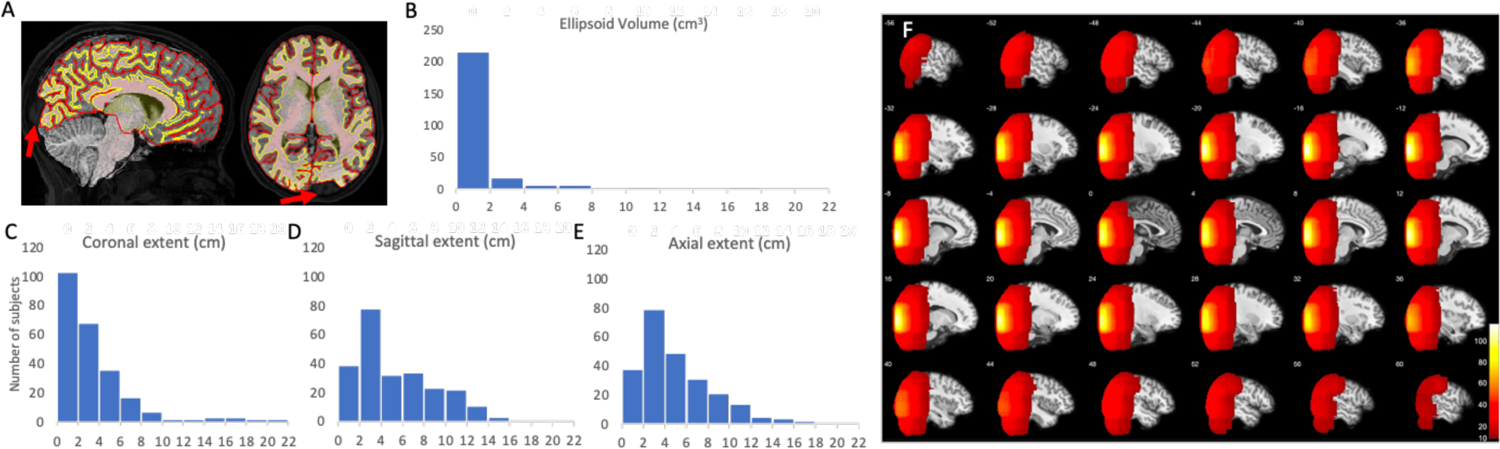
Signal dropout in sMRI processing (n=228). (A) Examples of dropout regions where FreeSurfer segmentation failed and did not include a substantial portion of cortex. (B) Distribution of approximate volume of dropout area estimated by ellipsoid volume calculated and distribution of (C) sagittal, (D) coronal, and (E) axial extent. (F) Distributions of drop-out regions overlaid on exemplar brain thresholded at n=10 subjects. Heat map represents number of overlapping subjects.

**Extended Figure 3.**
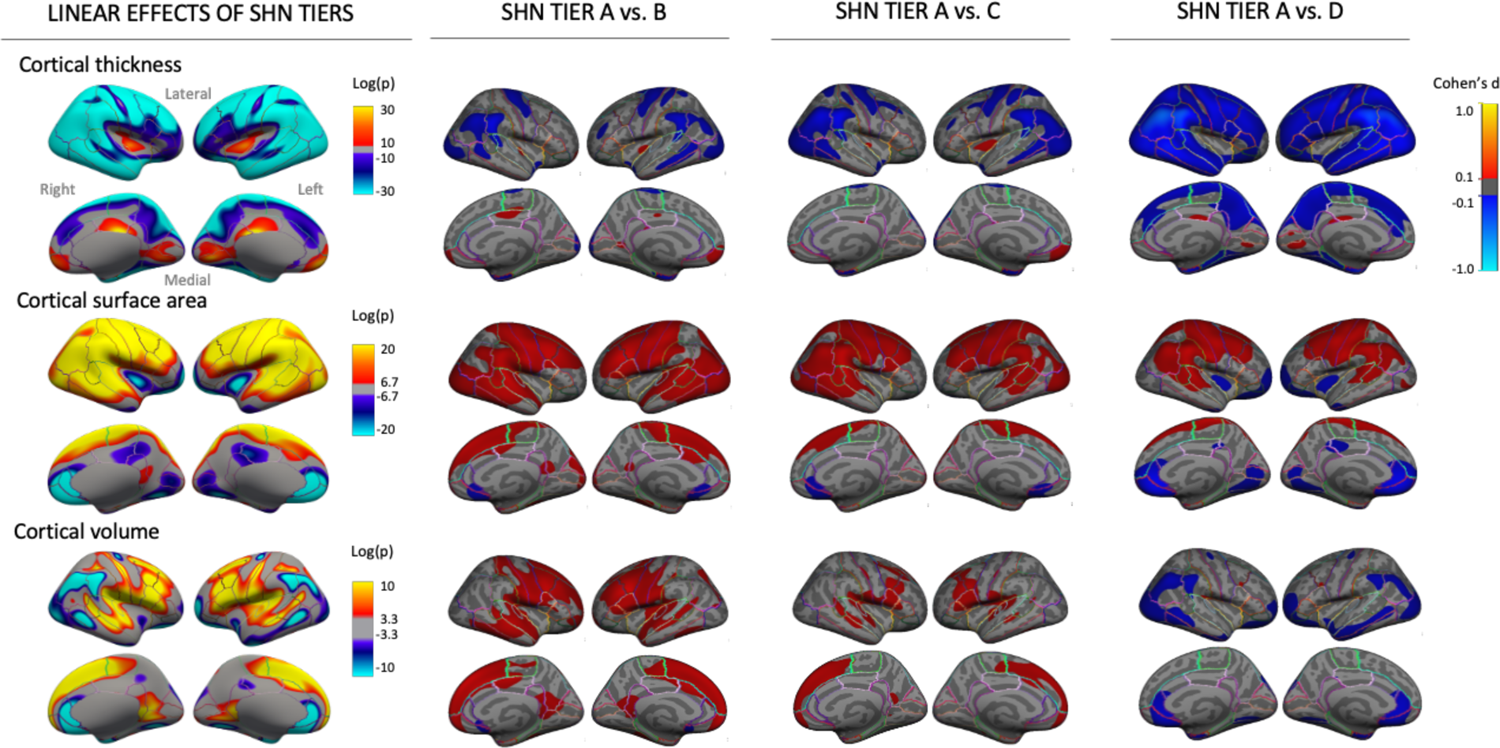
Comparison of SHN tier effects on sMRI indices at Baseline, n=10,295; compare to. **Figure 2.** Maps at left show linear associations of SHN tier (A to D) with cortical thickness, surface area, and volume. Maps at right contrast thickness, surface area, and volume highest quality images (SHN=A) with those assigned to lower quality ratings. Covariates included age, gender, estimated intracranial volume (fixed effects), site, and scanner manufacturer (random effects).

**Extended Data Figure 4.**
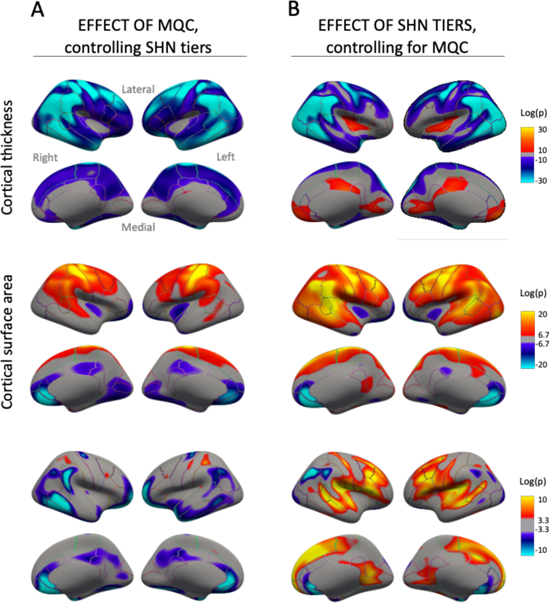
Unique contributions of SHN tiers versus MQC to variance in sMRI indices, n=10,295. (A) Linear association of MQC on cortical indices after controlling for SHN tiers. (B) Linear association of SHN tiers on cortical indices after controlling for MQC. Covariates included age, gender, estimated intracranial volume (fixed effects), site, and scanner manufacturer (random effects).

**Extended Data Figure 5.**
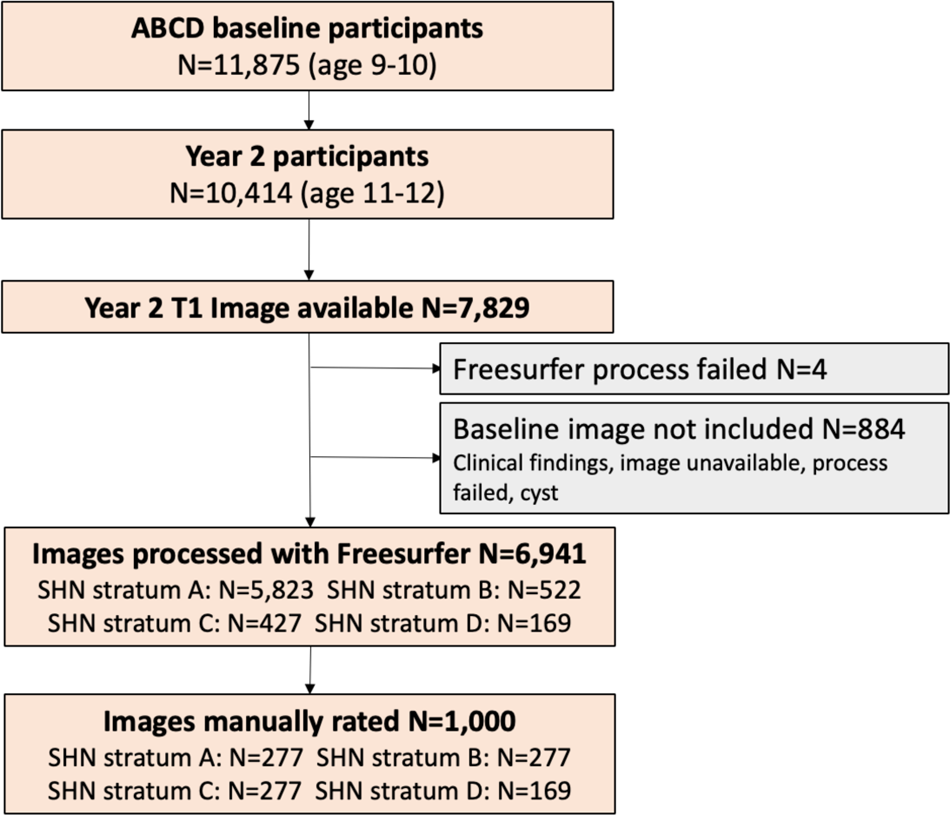
Included Year 2 follow-up scans. Among 11,875 total participants at baseline, Year 2 T1 scans were available from 7,829; of these, 6,941 were eligible for processing with FreeSurfer, and 1,000 were semi-randomly selected for MQC ratings (see Methods for additional details).

**Extended Data Figure 6.**
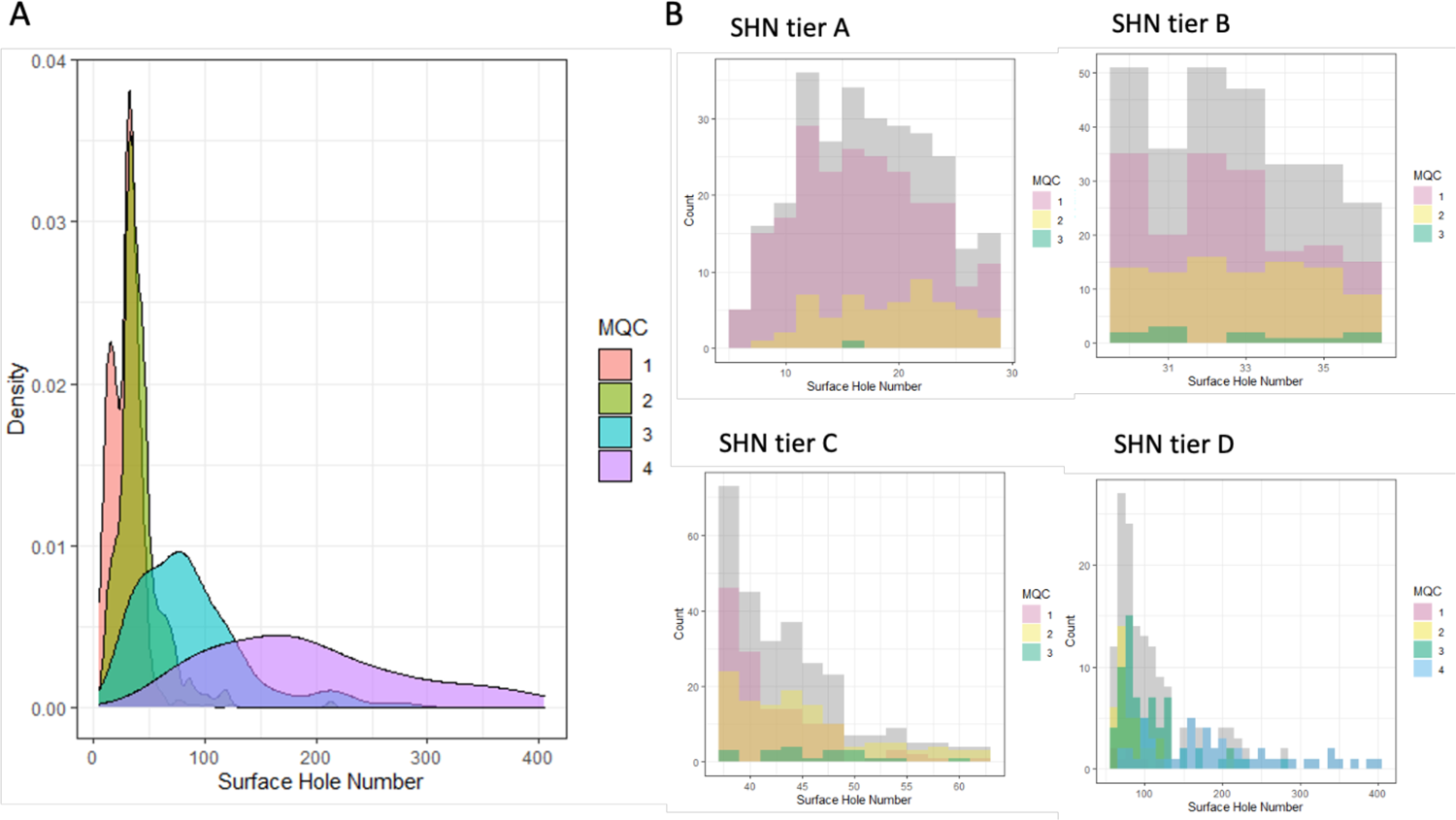
Relationship of surface hole number (SHN) to manual quality control (MQC) in selected Year 2 follow-up scans (n=999). (A) Density plot of SHN values, stratified by MQC ratings. (B) Distribution of MQC ratings as related to SHN for each SHN tier.

**Extended Figure 7.**
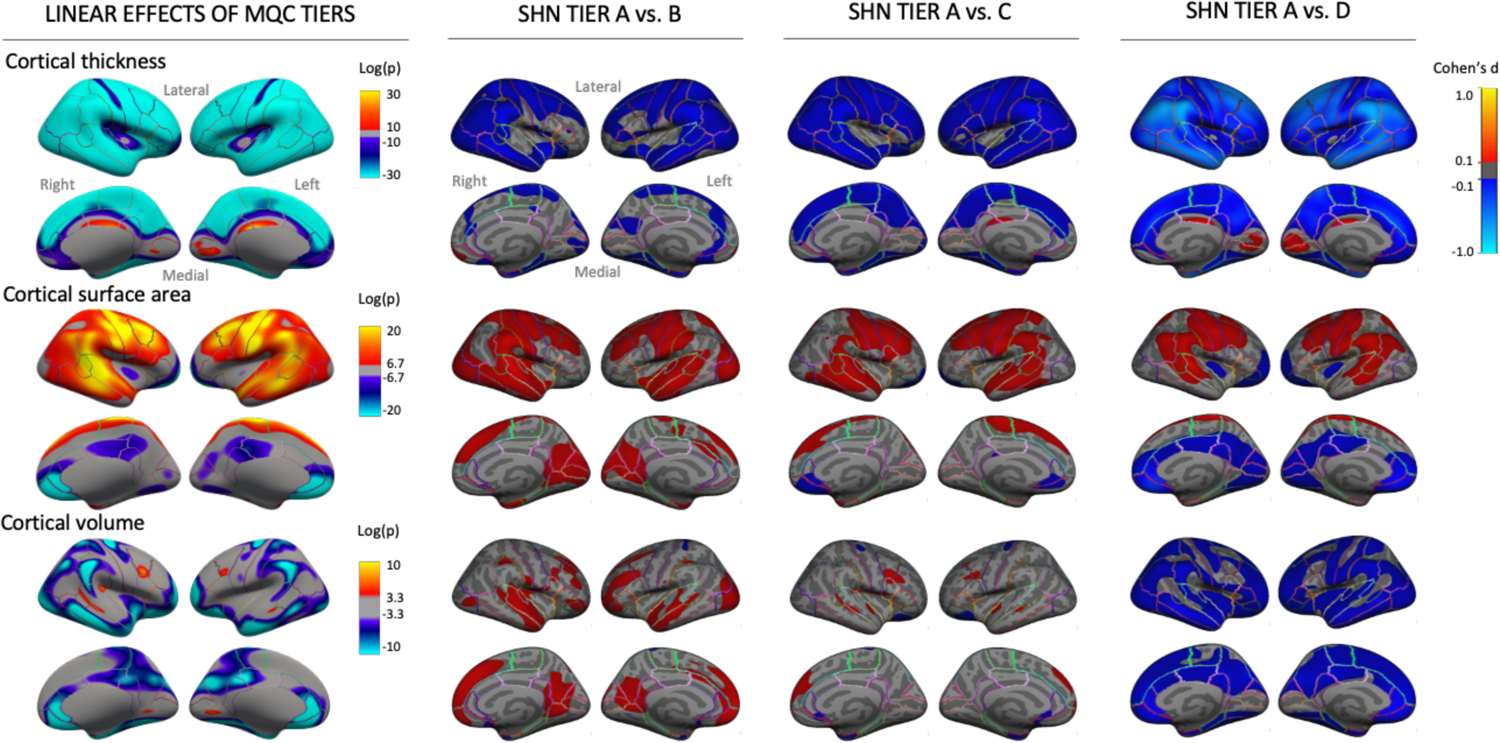
SHN tier effects on sMRI indices at Year 2, n=6,941 (compare to Extended Data Figure 3). Maps at left show linear associations of SHN tier (A to D) with cortical thickness, surface area, and volume. Maps at right contrast thickness, surface area, and volume highest quality images (SHN=A) with those assigned to lower quality ratings. Covariates included age, gender, estimated intracranial volume (fixed effects), site, and scanner manufacturer (random effects).

**Extended Data Figure 8.**
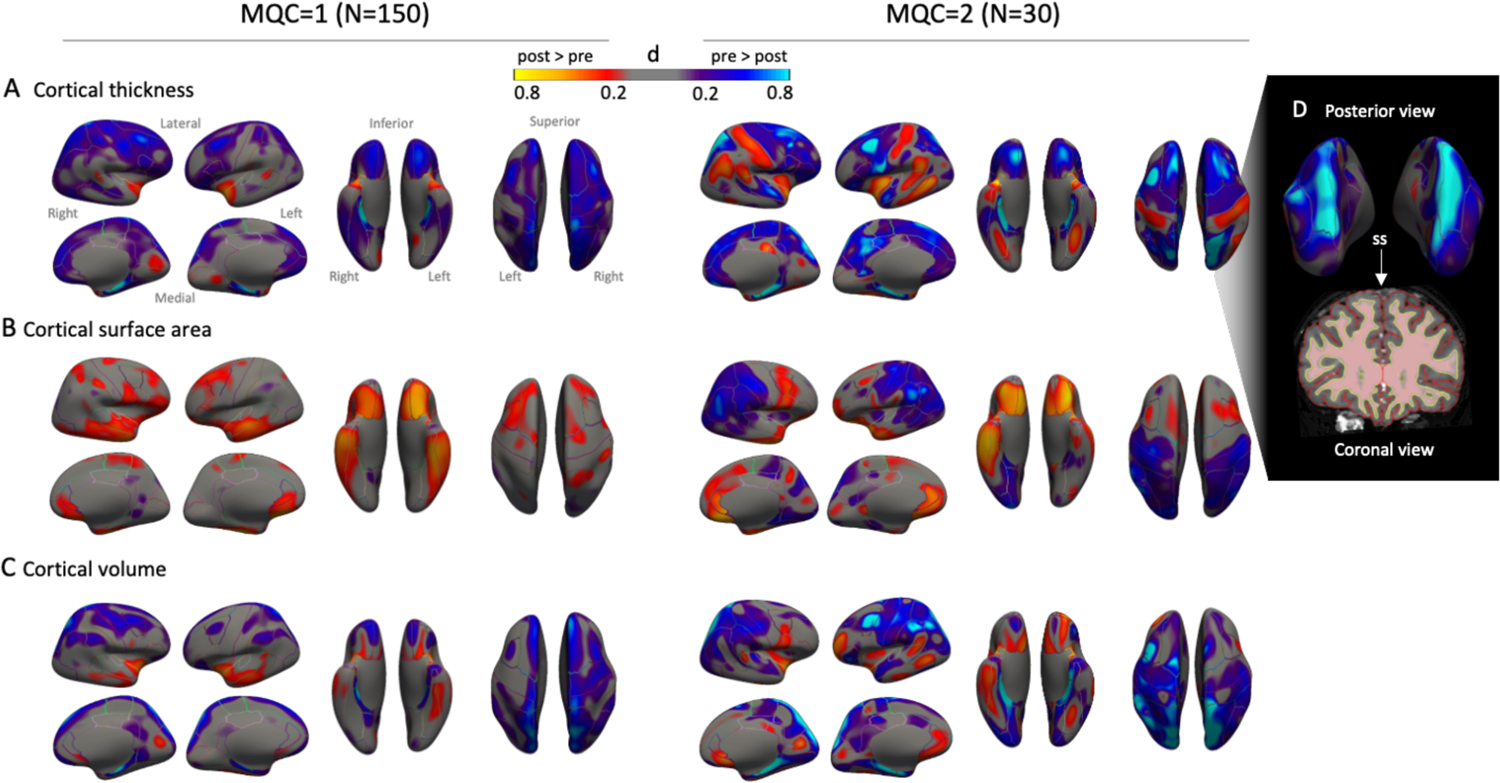
Effects of manual edits on sMRI indices, stratified by MQC rating. Edits were conducted on 150 scans with MQC=1 and 30 scans with MQC=2. Maps reflect effect sizes of pre-to-post edit changes in (A) cortical thickness, (B) cortical surface area, and (C) cortical volume. Note increased effects of edits in MQC=2 relative to MQC=1. (D) Post-edit thickness reduction along the superior sagittal sinus, which is frequently misattributed to pial surface during preprocessing.

**Extended Data Figure 9.**
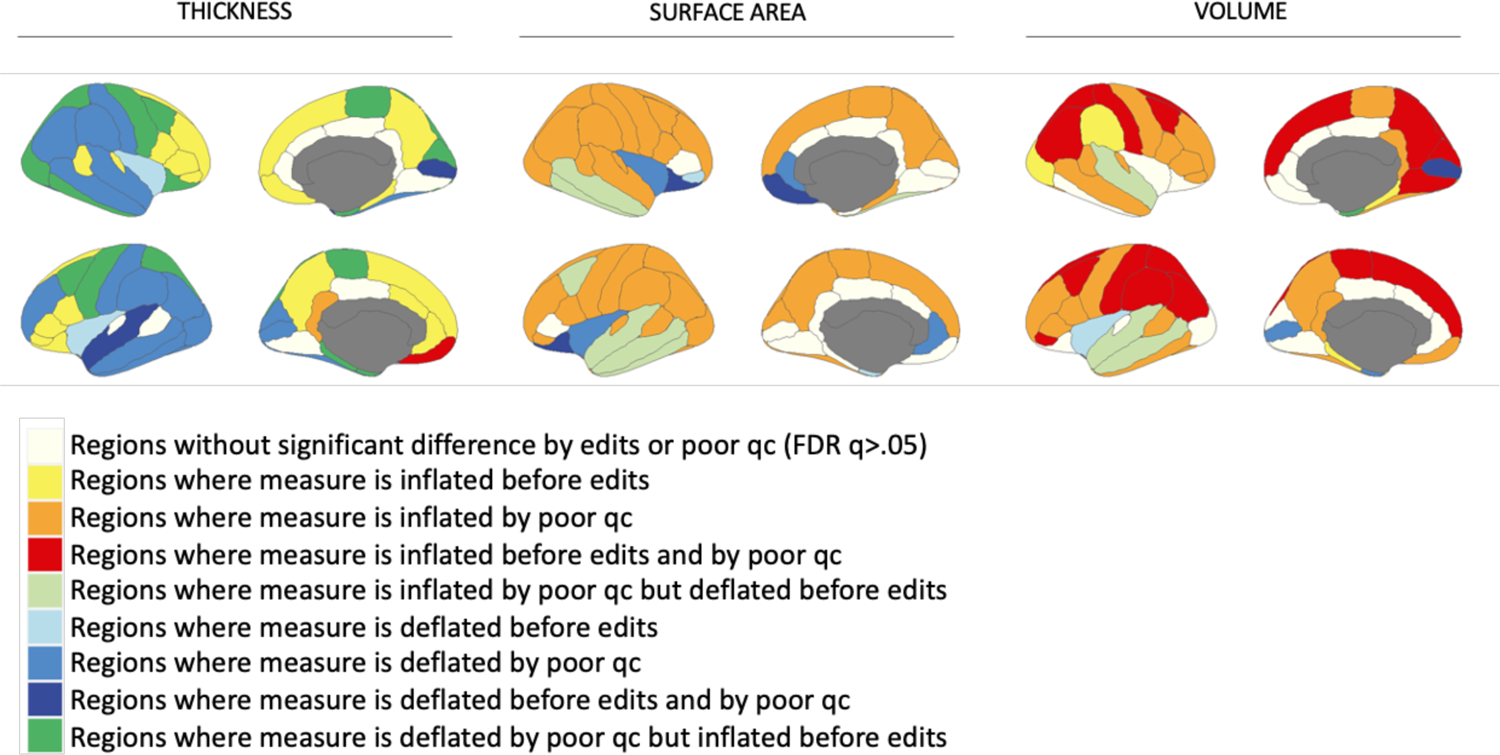
Composite maps showing location and direction of sMRI measurement errors detected by manual quality control and cortical edits, among MQC=1 and 2 scans only. Highlighted regions show either significant differences in sMRI indices between MQC=1 and MQC=2 scans, significant effects of cortical edits, or both. Note that, when co-occurring within the same region, errors due to poor scan quality (assessed by MQC) do not necessarily occur in the same direction as errors requiring manual edits.

**Extended Data Figure 10.**
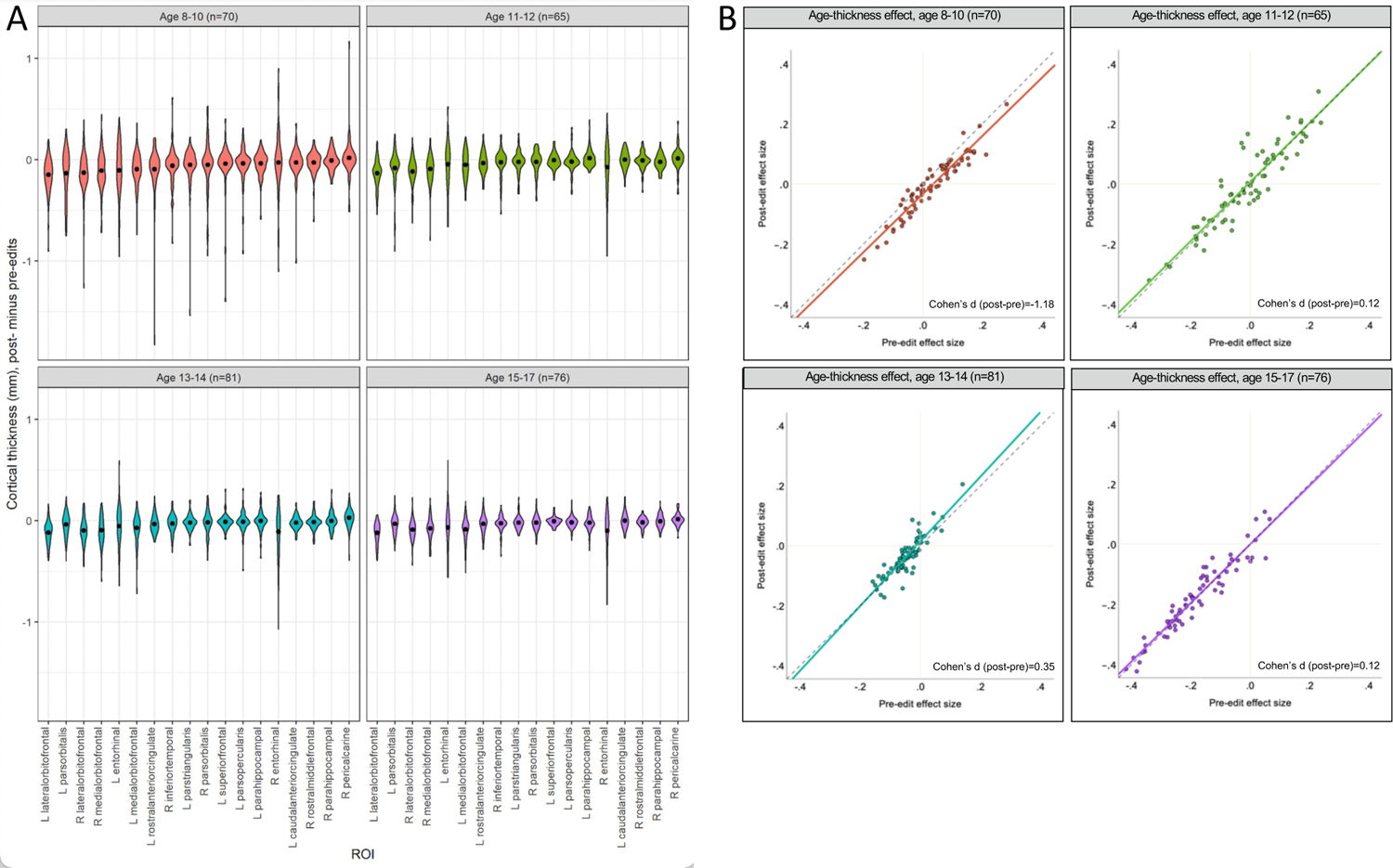
Effects of manual edits on cortical thickness and age-thickness relationships MGH sample, stratified by age group (n=292). (A) Violin plots show effect size and related variance of manual edits on cortical thickness in the MGH sample, stratified by age group. The 18 included ROIs are those that also showed significant effects of edits on cortical thickness in the ABCD cohort, in the same direction. Regions are ordered by effect size in the 8- to 10-year-old group. Means are represented by black circles. Note that effect sizes and variance diminished with age. (B) Effects of edits on the magnitude of age-thickness relationships within the MGH sample across 68 cortical ROIs, stratified by age group. Each marker shows the age-thickness effect size for a given ROI. Edits strengthened age-thickness effects (i.e., effect sizes became more negative, indicated by lower intercept of the best-fit line compared to the dashed unity line) at age 8-10, but not in other age groups.

## Acknowledgments

The authors are grateful to Drs. Randy L. Buckner and Erin C. Dunn for helpful comments on the manuscript.

## Author contributions

**Conception and experimental design:** Kunitoki, Clauss, Doyle, Lee, Tervo-Clemmens, Eryilmaz, Satterthwaite, Roffman.

**Data acquisition:** Hopkinson, Eryilmaz, Gollub, Dowling, Roffman.

**Data analysis:** Elyounssi, Kunitoki, Clauss, Laurent, Kane, Hughes, Bazer, Sussman, Lee, Dowling, Roffman.

**Data interpretation:** Elyounssi, Kunitoki, Class, Laurent, Kane, Hughes, Bazer, Doyle, Lee, Tervo-Clemmens, Gollub, Barch, Satterthwaite, Dowling, Roffman.

**Drafting and revision of manuscript:** All authors. All authors have approved the submitted version of the manuscript and have agreed to be personally accountable to their own contributions.

